# Quantifying cell line specific proliferation and migration rates in glioblastoma cells

**DOI:** 10.1101/654053

**Authors:** Emil Rosén, Philip Gerlee, Sven Nelander

**Affiliations:** Department of Immunology, Genetics and Pathology Uppsala University; Mathematical Sciences Chalmers University of Technology & University of Gothenburg

## Abstract

We have characterised the migration and proliferation rates of a large number of patient-derived glioblastoma cell lines using an individual-based model coupled to an Approximate Bayesian Computation algorithm. We found that the cell lines exhibited a negative correlation between the rate of migration and rate of division. This observation agrees with the Go or Grow hypothesis and highlights patient-specific differences in migration and proliferation.

## I. Introduction

The brain tumour glioblastoma grade IV (GBM) is characterised by a short median survival of 14 months after diagnosis [4]. This is in part due to diffuse growth patterns driven by high rates of cellular migration. Thus, inhibition of migration pathways might constitute an interesting complement to standard glioblastoma therapies that seek to inhibit cell proliferation rate. However, the potential of migration as a therapeutic target is complicated by the strong dependency between migration and proliferation phenotypes, which is known as the Go or Grow hypothesis [2]. Our aim here is to characterise migration and proliferation rates in panel of 69 patient derived cell lines with the future aim of assessing drug effects on these traits in a patient-specific manner.

## II. Results

The data that we have analysed was obtained from a large-scale *in vitro* drug screen primarily aimed at finding drugs that reduced cell division in GBM cells. The data acquired was a single microscopy image taken after 72 hours (see fig. 1 and Methods). A simple cell count of the microscopy image is informative about the effect of the drug on cell division, but is it possible to also extract information about potential effects on cell migration? This is indeed possible and the reason is the fingerprint left by cell division and migration on the spatial correlations in the image. The process of cell division, which leaves two daughter cells in close proximity, increases spatial correlations at short distances, while cell migration, which is random and unbiased in *in vitro* conditions, tends to separate daughter cells and hence reduce spatial correlations.

**Fig. 1.**
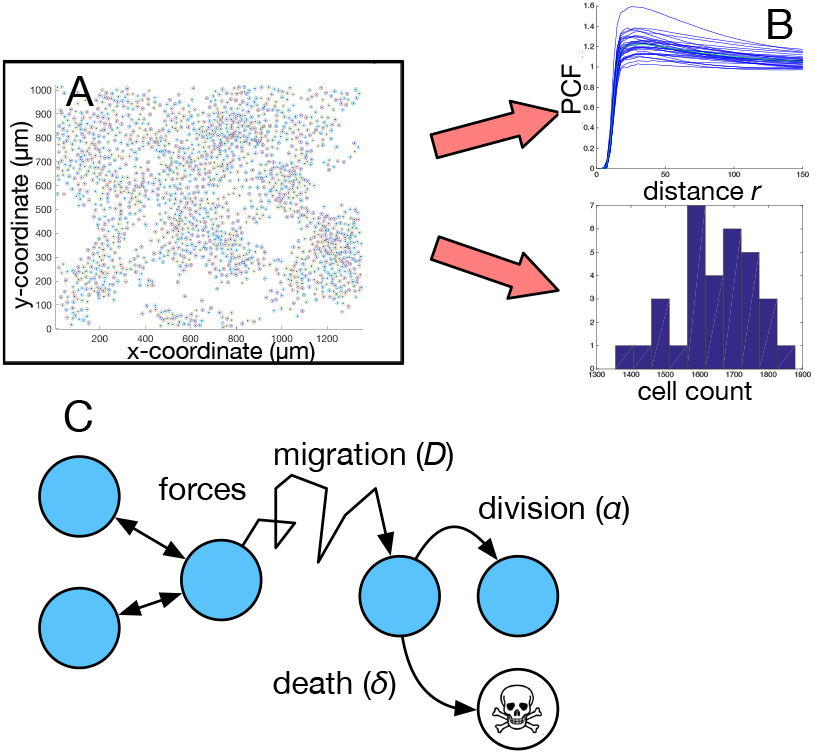
Overview of data analysis and model. (A) After 72 hours of growth the position of the GBM-cells in each well were recorded. (B) From this data the pair correlation function (PCF) and the number of cells was calculated. (C) In the off-lattice individual-based model the cells interact mechanically, migrate, divide and die.

In order to quantify this effect, we calculated the pair correlation function (PCF) [3], which measures the normalised density of cell pairs separated by distance *r* (see fig. 1). We found a large variation in the PCF for different cell lines even under control conditions suggesting that the spatial arrangement (and possibly the migration and division rates) of cells in the images varied in a cell line specific manner.

To get a deeper understanding of the connection between rates of cell migration and proliferation and the cell count and PCF we constructed an individual-based model (IBM) of the *in vitro* assay. In this model the GBM cells are represented as particles in a 2-dimensional domain that interact mechanically, migrate, divide and die (see fig. 1 and Methods). In order to fit the model to the data we used an Approximate Bayesian Computation algorithm that used both the cell count and the PCF to infer the model parameters (see Methods) [3].

We found that the proliferation rates and diffusion coefficients varied across the 69 cell lines. For a subset of cell lines it was not possible to get reliable estimates of the diffusion coefficient, and these were excluded from the analysis. Within the remaining cell lines there was a notable positive correlation between diffusion coefficient and doubling time (see fig. 2).

**Fig. 2.**
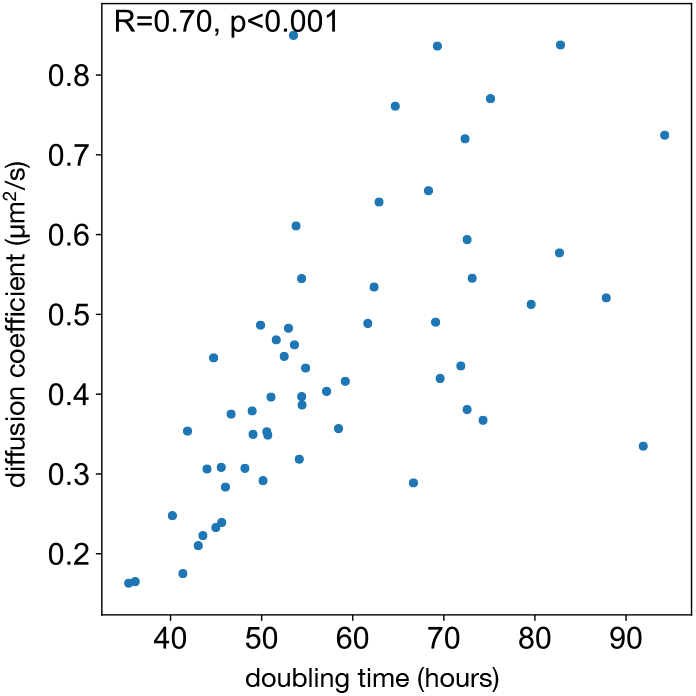
Scatter plot of the estimated doubling times and diffusion coefficients of the GBM cell lines. Each point corresponds to a cell line and we observe a strong positive correlation (Spearman’s rank correlation *ρ* = 0.70) between diffusion coefficient and doubling time.

## III. Discussion

Our results lends support to the long-standing Go or Grow hypothesis. It should be noted that we have quantified migration rates on laminin coated plates, and the extent to which this correlates to *in vitro* migration remains to be elucidated. Nevertheless, we observe a trade-off between migration and proliferation which highlights that cytoskeleton dynamics could be limiting, as it is involved in both cell division and force generation in migration.

The parameter estimates obtained from a single static image will be validated by performing time-lapse microscopy on selected number of cell lines. From images taken at 30 minute intervals it will be possible to get more reliable estimates of diffusion and cell division rates, possibly at the level of single cells. The correlations to *in vitro* behaviour will be assessed by comparing the inferred parameters to existing data from an orthotopic mouse model. We will also compare predicted tumour growth dynamics to MRI-data for patients from which the cell lines were derived. In the future we will also analyse images obtained under drug treatments paving the way for patient-specific mapping of drug effects on GBM migration and proliferation.

## IV. Methods

### Data acquisition

69 cell lines from the Human Glioma Cell Culture (HGCC) resource [5] were suspended in stem cell medium and plated on 384 well plates (BD Falcon Optilux #353962) coated in laminin at a density of 1000-2200 cells/well. The cells were cultured at 37 °C and 5 % CO_2_ for 72 hours and imaged using a Operetta and ImageXpress microscope at 20x magnification. The images were segmented and the position of the cells were recorded. For each well we counted the total number of cells and calculated the pair correlation function.

### Individual-based model

We model the well in which the cells migrate, divide and die as a square region in two-dimensional space with side length *L*. The position of cell *i* is denoted *x*_*i*_(*t*) and to denote the position of all cells we write 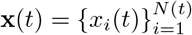, where *N* (*t*) is the number of cells at time *t*.

We assume that cell motion is over-damped and therefore model cell migration and mechanical interactions using a Langevin equation [1]: 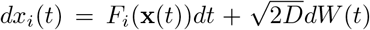, where migration is modelled as a Brownian motion with diffusion coefficient *D* and inter-cellular forces (pushing/adhesion) are captured by the drift term *F*_*i*_(**x**(*t*)), which depends on the position of all other cells. The total force is assumed to be a sum of pairwise forces, which are non-zero only if the cells are within a distance *R* (the cell radius) and hence overlap.

We assume that cell division occurs with a base line rate *α* > 0, and is reduced due to contact inhibition by the presence of other cells. This is modelled using an interaction kernel *w*_*b*_(*r*) = *e*^−*γr*^, which only depends on the distance *r* between the cells. The total rate of cell division for cell *i* with position *x*_*i*_ is given by 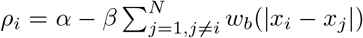 Upon cell division the daughter cell is placed at distance *R* from the parent cell, and at an angle drawn uniformly from the interval [0, 2*π*]. Lastly, we assume that cell death occurs with rate *δ* independent of cell density.

The system is initialised by placing *N*_0_ cells at random in the domain. To simulate the system we use a combination of the the Gillespie algorithm (for discrete events) and the Euler– Maruyama method (for cell migration). The simulations were carried out for 72 hours, and the number of cells and the PCF was calculated.

In order to find the model parameters that best describe the data we used Approximate Bayesian Computation (ABC). As summary statistics we used an equally weighted sum of the cell count and the pair correlation function (see [3] for details). The sparse data made it difficult to infer death rates and we therefore set *δ* = 0 and instead absorbed cell death into the baseline proliferation rate.

